# Circadian dynamics of the Zbtb14 protein in the ventral hippocampus are disrupted in epileptic mice

**DOI:** 10.1101/2024.03.07.583828

**Authors:** İlke Güntan, Antoine Ghestem, Kinga Nazaruk, Karolina Nizińska, Maciej Olszewski, Dorota Nowicka, Christophe Bernard, Katarzyna Łukasiuk

## Abstract

Our previous *in silico* data indicated an overrepresentation of the ZF5 motif in the promoters of genes in which circadian oscillations are altered in the ventral hippocampus in the pilocarpine model of temporal lobe epilepsy in mice. In this study, we test the hypothesis that the Zbtb14 protein oscillates in the hippocampus in a circadian manner and that this oscillation is disrupted by epilepsy.

We found that Zbtb14 immunostaining is present in the cytoplasm and cell nuclei. Western blot data indicate that the cytoplasmic and nuclear levels of Zbtb14 protein oscillate, but the phase is shifted. The densities of the Zbtb14-immunopositive express circadian dynamics in the ventral hilus and Ca3 but not in the dorsal hilus, Ca3, or the somatosensory cortex. In the pilocarpine model of epilepsy increase in the level of Zbtb14 protein was found at 11 PM, but not at 3 PM compared to controls. Finally, *in silico* analysis revealed the presence of the ZF5 motif in the promoters of 21 out of 24 genes down-regulated by epileptiform discharges *in vitro*, many of which are involved in neuronal plasticity. Our data suggest that Zbtb14 may be involved in the circadian dynamic of seizure regulation or brain response to seizure rhythmicity.

**Highlights:**

- the Zbtb14 protein is expressed in neurons in the mouse brain;
- Zbtb14 protein levels oscillate through the circadian cycle in the ventral hippocampus but not in the dorsal hippocampus;
- the oscillations of the Zbtb14 protein occur in both the cytoplasm and nucleus but in a different temporal pattern;
- the circadian dynamics of the Zbtb14 protein are perturbed in epilepsy in an *in vivo* model of epilepsy;
- numerous genes that are downregulated in the *in vitro* model of epileptiform discharges have a ZF5 motif in their promoters

## Introduction

The circadian rhythm is a biological oscillation of around a 24-hour period, which is mostly regulated by the suprachiasmatic nucleus (SCN). This inner clock controls essential functions like the sleep-wake cycle, body temperature, hormone secretion, blood pressure, and immune functions (Hetman et al., 2022; Markov and Goldman, 2006). Although mammals have a self-regulated endogenous clock, this body clock needs to be adjusted to 24 hours by the external cues called zeitgebers. Peripheral organs transmit these external cues to the SCN of the hypothalamus, and SCN neurons coordinate the peripheral clocks (Hetman et al., 2022; Schurhoff and Toborek, 2023).

Self-sustaining transcriptional and translational feedback loops within SCN and other tissues synchronize the self-regulated endogenous clock and external cues. The cycle starts with the heterodimerization of brain muscle aryl hydrocarbon receptor nuclear translocator-like 1 (BMAL1; also known as ARNTL) and circadian locomotor output cycle kaput (CLOCK) - the positive regulators, which, in turn, activate the expression of its negative regulators, cryptochrome-1/2 (CRY1,2) and period-1/2/3 (PER1,2,3), among other downstream circadian rhythm modulators like RORα, REV-ERBα, and REV-ERBβ (Hastings et al., 2018).

Circadian rhythms are modified in numerous neurological disorders (Hetman et al., 2022; Lane et al., 2023; Rijo-Ferreira and Takahashi, 2019), including epilepsy (Debski et al., 2020). Conversely, disrupting the circadian machinery may lead to epilepsy (Li et al., 2017). Most, if not all, patients with epilepsy present a circadian regulation of their seizures (Karoly et al., 2021), a property also observed in experimental models (Baud et al., 2019; Quigg et al., 1998). This oscillatory pattern of seizures may be linked to altered circadian rhythms (Bernard, 2021).

In epileptogenic regions, like the hippocampus, the circadian rhythmicity is altered. The expression of seven core genes (BMAL1, CLOCK, CRY1/2, and PER1/2/3) is perturbed in the hippocampus in an experimental model of temporal lobe epilepsy (Matos et al., 2018). We have shown that in both control experimental epilepsy conditions, the levels of a large number of transcripts and proteins oscillate in a circadian manner (Debski et al., 2020). However, the pattern is modified, manifested by phase-shift and amplitude changes of many transcripts (Debski et al., 2020). The origin of these alterations is not known. Interestingly, promoter analysis revealed the overrepresentation of the ZF5 motif in groups of genes representing specific expression perturbations in epilepsy (Debski et al., 2020). We thus hypothesized that proteins with ZF5 binding motifs should also oscillate to induce oscillations of their targets.

The Zbtb14 protein (formerly also known as ZNF478, ZFP161, ZFP5, ZF5) binds to ZF5 motif on gene promoters and is expressed in the brain (https://www.genecards.org/cgi-bin/carddisp.pl?gene=ZBTB14) (Obata et al., 1999; Yokoro et al., 1998). Zbtb14 protein is poorly characterized. It contains five C2H2 (Krüppel-type) zinc fingers on its C-terminus and POZ/BTB (Broad-complex, Tramtrack and Bric-a-brac/Poxvirus and Zinc-finger) domain in its N-terminus (Numoto et al., 1993). It is expressed in several human and mouse tissues, including the testis, liver, brain, muscle, thymus, and spleen (Yokoro et al., 1998). Zbtb14 can act as a transcriptional repressor, e.g., of the human fragile X-mental retardation 1 (FMR1) gene and leukemia inhibitory factor (Lif), or as a transcriptional activator, e.g., of the human dopamine transporter (hDAT) and interleukin 6 (Il6) (Lee et al., 2004).

In this study, we test the hypothesis that the Zbtb14 protein oscillates in the hippocampus in a circadian manner and that this oscillation is disrupted by epilepsy.

## Materials and Methods

### Animals, time-pointed tissue collection, and epilepsy induction

All animal procedures were conducted on FVB mice following European Council Directive 2010/63/EU and ARRIVE guidelines.

Naïve animals for time-pointed tissue collection are obtained from the Nencki Institute Animal Facility and performed at the Nencki Institute of Experimental Biology (Polish Academy of Sciences, Warsaw, Poland). All experiments on control and epileptic animals were performed at INSERM, INS, Institut de Neurosciences des Systèmes, Aix Marseille University, Marseille, France, following INSERM procedures on FVB adult male mice as described in Debski et al. (Debski et al., 2020).

Mice were housed in a controlled environment (7:30 am-7:30 pm, light on/off) with water and food available *ad libitum*. During the dark phase, a red light was used while handling animals. Any other light source was off to ensure the circadian rhythm was unaffected. The animals were kept together in groups (max. six animals per cage), and cardboard tunnels and snacks enriched their environment. In addition, the same researchers took care of the animals to reduce the external stressors.

For time-pointed tissue collection, animals aged of 16th and 19th weeks were anesthetized with isoflurane at 11 am, 3 pm, 7 pm, 11 pm, 3 am, and 7 am. The brain’s left hemispheres were fresh-frozen on dry ice for immunofluorescent staining and stored at -80°C. The right hippocampi were removed from the hemispheres and immediately used to extract nuclear and cytoplasmic protein extracts.

To induce epilepsy, adult FVB mice were injected with methylscopolamine [1 mg/kg, intraperitoneally (ip)] 30 min before the pilocarpine injections. After that, pilocarpine was repeatedly injected (100 mg/kg, ip) every 20 minutes until status epilepticus (SE) was observed. After 90 minutes of SE, we injected diazepam (10 mg/kg, ip) to stop SE. All mice then received 0.5 ml of NaCl (0.9%) subcutaneously and again in the evening. If required, mice were fed using a syringe during the following days. Control mice had the same treatment but were only injected with intraperitoneal NaCl (0.9%). Animals were anesthetized with isoflurane in the animal facility and were sacrificed at the same time points: 3 pm and 11 pm. The hippocampus was removed in modified ice-cold artificial cerebrospinal fluid (ACSF). Both hippocampi from the same animal were quickly frozen and stored together at -80°C.

### Immunofluorescent double staining

Sagittal sections (20 μm) were cut with a cryostat (Leica CM1860), collected on poly-L-lysine-coated glass microscope slides, and stored at -80°C. Sections were fixed in ice-cold acetone for 10 minutes. Slides were washed in 0.5% Triton X-100 (Sigma) in PBS (PBST) thrice for 5 minutes. Unspecific binding was blocked by 90 minutes of incubation with 5% normal goat serum (Vector, #S-1000) at room temperature. Next, the sections were incubated overnight in a humid chamber with rabbit anti-ZBTB14 antibody (1:100; Atlas Antibodies, #HPA050758) diluted in PBST at 4°C. The following day, the slides were washed in PBST three times for 5 minutes and incubated for 2 hours with goat anti-rabbit IgG antibody (H+L), biotinylated (1:200; Vector, BA-1000) diluted in PBST at room temperature. This is followed by washing and incubation for 20 minutes with fluorescein avidin D, FITC (1:100; Vector, A-2001) at room temperature. After washing, the sections were blocked at room temperature for 90 minutes of incubation with 2% normal donkey serum (Jackson ImmunoResearch, #017-000-121). Next, the sections were incubated overnight in a humid chamber with mouse anti-NeuN antibody, clone A60 (1:1000; Sigma-Aldrich, #MAB377) diluted in PBST at 4°C. The following day, the slides were washed and incubated for 2 hours with donkey anti-mouse IgG (H+L) highly cross-adsorbed secondary antibody, Alexa Fluor 568 (1:2000; Invitrogen, #A-10037) diluted in PBST at room temperature. Nuclei were stained with Hoechst dye (1:2000; Sigma-Aldrich). Finally, the slides were washed in PBS thrice for 5 minutes and rinsed in cold tap water. They were dried overnight and cover-slipped with Vectashield Mounting Medium (Vector, #H-1000) the next day, stored at 4°C. The omission of primary antibodies verified the specificity of antibodies.

### Quantification of the density of Zbtb14-positive cells

Sections were viewed with a Nikon Eclipse 80i microscope, and images were captured using a CCD camera (Evolution VF; MediaCybernetics) with a Nikon 10/0.30 DIC L/N1 or 20/0.50 DIC M/N2 objective. Images were acquired for each area examined using different filters for Alexa Flour 568, FITC, and Hoechst and elaborated with Image-ProPlus, version 5.0 for Windows (MediaCybernetics). The sections from the 2.28 – 2.40 mm lateral level from the midline were visually inspected under the microscope. All images were collected using the same exposure time, and then corresponding pictures were superimposed to visualize the co-localization of the NeuN-stained neurons with the Zbtb14 immunofluorescence. Data were collected from a manually delineated area covering the hilus (separately ventral and dorsal areas), CA3 area (averaged from two separate frames covering CA3a), and somatosensory cortex (−2 from bregma, same sections as for hilus). The hilus was distinguished according to Amaral et al. (Amaral et al., 2007). For the cortex, data was calculated from a 200 μm-wide profile of the somatosensory cortex. The profile was divided into cortical layers II/III, IV, V, and VI according to Hoechst staining, which was evaluated separately. The Zbtb14-positive cells were manually tagged and counted within the area of interest. NeuN/Zbtb14 double-stained cells were also recorded in the ventral hilus. Only the cells with well-visible nucleoli (visualized with Hoechst staining) were included in the analysis. The expression for each marker was analyzed as the number of positive cells per mm^2^. Figures were prepared using Adobe Photoshop CS2 and Corel Draw X4. Brightness and contrast were adjusted to regain the natural appearance of the sections.

### Protein extraction and western blot

The right hippocampi from the time-pointed tissue collection are used for nuclear and cytoplasmic protein extraction. The nuclear and cytoplasmic protein extraction was performed using NE-PER™ Nuclear and Cytoplasmic Extraction Reagents kit (Thermo Scientific, #78833) according to the manufacturer’s instructions. The concentration of the protein extracts was measured using Protein Assay Dye Reagent (BioRad, #5000006).

The fresh-frozen brain tissue from control and epileptic animals was homogenized with a handheld rotor-stator homogenizer (QIAGEN, TissueRuptor II) in a lysis buffer that contains 0.5% Triton X-100, 50 mM potassium chloride, 50 mM PIPES, 2 mM magnesium chloride, and 20 mM EGTA with the addition of 0.1 mM phenylmethylsulfonyl fluoride, 1 mM dithiothreitol, 1X proteinase inhibitor cocktail (Roche, #11697498001). The tissue lysate was frozen for 20 minutes at -20°C. After thawing, the samples were centrifuged at 4°C, 11000 rpm, for 20 minutes; the supernatant was transferred into a clean tube and stored at -80°C. The concentration of the protein isolates was measured using Protein Assay Dye Reagent (BioRad, #5000006). 20 μg of protein for cytoplasmic extracts from naïve animals, control and epileptic animal total protein isolates, and 4 μg nuclear protein samples were loaded for Western Blot. Protein isolates were separated via Tris-glycine sodium dodecyl sulfate-polyacrylamide gel electrophoresis (SDS-PAGE) and transferred to a nitrocellulose blotting membrane (Cytiva, #10600002).

The membranes were blocked in 5% nonfat milk in TBS-T (150 mM NaCl, 10 mM Tris, and 0.1% Tween20, pH 8.0) for 1 hour at room temperature. Next, they were incubated overnight in rabbit anti-ZBTB14 (1:1000; Atlas Antibodies, #HPA050758) diluted in TBS-T at 4°C. After washing with TBS-T, the membranes were incubated for 2 hours in goat anti-rabbit IgG antibody, peroxidase-conjugated (1:10000; Sigma-Aldrich, #AP132P) diluted in TBS-T at room temperature. The signal was detected using ECL Prime Western Blotting System (Cytiva, #RPN2232) according to the manufacturer’s instructions. The signal was captured on an X-ray film with an automatic film processor. After stripping (93 mM glycine, 69 mM sodium dodecyl sulfate, pH 3.0), the membranes were re-probed with anti-β-actin−peroxidase antibody, mouse monoclonal (1:240000; Sigma-Aldrich, #A3854) diluted in TBS-T. X-ray films were scanned using GS-900 Calibrated Densitometer (BioRad), and optical density was measured using Image Lab Software version 5.2.1. GraphPad Prism version 5.01 was used for statistical analysis.

### In vitro model of epileptiform discharges

Embryos at 18 days post-fertilization (E18) were used to establish primary cultures of cortical neurons (Xu et al., 2012). Isolated embryos were decapitated, the hemispheres were isolated in a chilled HBSS buffer (Thermo Fisher, #14170-088), and incubated at 37°C for 15 minutes with HBSS buffer containing 0.2% Trypsin (Thermo Fisher, #27250-0180) and 0.15 mg/ml DNase (Sigma-Aldrich, #DN-25). The solution was removed, and warm 10% FBS (Thermo Fisher, #10106-151) diluted in HBSS was added. After washing the tissue twice with fresh, warm HBSS without FBS, 2 ml of warm medium containing 1x B-27, 10% FBS, 10 mg/ml Gentamicin, 0.5 mM Glutamax in Neurobasal Medium was added, and then the tissue was pipetted several times. The number and viability of cells were measured in 0.4% Trypan Blue using a Neubauer chamber (Marienfeld). Two hundred thousand cells per well were seeded into a 12-well plate pre-coated with poly-D-lysine (5 μg/ml) in 0.1 M borate buffer (Sigma-Aldrich, #P7280). Cultures were grown in an incubator at 37°C and 5% CO_2_. Half of the medium was exchanged for FBS serum-free medium (1x B-27, 10 mg/ml Gentamicin, 0.5 Mm Glutamax in Neurobasal Medium) on days 2 and 6.

Epileptiform discharges were induced *in vitro* as described by Jiang et al. (Jiang et al., 2010) with modifications (Nizińska K et al. submitted). Briefly, cultures were incubated for 3 hours in pBRS buffer without magnesium (145 mM NaCl, 2.5 mM KCl, 10 mM HEPES, 2 mM CaCl2, 10 mM glucose, 0.002 mM glycine, pH=7.3). The control cultures were incubated in pBRS buffer with magnesium (145 mM NaCl, 2.5 mM KCl, 10 mM HEPES, 2 mM CaCl2, 10 mM glucose, 0.002 mM glycine, 1mM MgCl2, pH=7.3). At the end of the incubation, the pBRS buffer was replaced with a fresh, warmed, serum-free culture medium (37°C). Material for RNA-seq studies was collected 24 hours after induction of epileptiform discharges.

### RNA isolation and new-generation sequencing

RNA using Qiazole (Qiagen, #79306) and RNeasy Mini Kit (Qiagen, #74104) according to the manufacturer’s instructions. The RNA was measured on a Nanodrop Spectrophotometer (DeNovix, #DS-11 Spectrophotometer) at λ= 260 nm and 280 nm. The isolated RNA was stored at -80°C. RNAseq libraries were prepared by KAPA Stranded mRNA Sample Preparation Kit according to the manufacturer’s protocol (Kapa Biosystems, MA, USA) as previously described (Grabowska et al., 2022). Transcriptomic data analysis was done as follows: fastq files were aligned to the rn6 rat reference genome with the STAR program (Dobin et al., 2013), and reads were counted to genes using the feature Counts algorithm (Liao et al., 2014). Gene counts were normalized with the FPKM method, and differential analysis was performed by DESeq2 (Love et al., 2014). Genes were considered to be differentially expressed with adjusted *p*-value < 0.05.

Analysis of the overrepresentation of transcription factor binding motifs in groups of genes was performed using g:Profiler (https://biit.cs.ut.ee/gprofiler/gost).

Data are deposited at the NCBI GEO repository with the submission number: GSE227084 (https://www.ncbi.nlm.nih.gov/geo/query/acc.cgi?acc=GSE227084).

## Results

### Localization of Zbtb14 immunostaining

Cellular localization of the Zbtb14 protein in the ventral hippocampus and somatosensory cortex of control mice was evaluated by immunohistochemistry. We focus on the ventral hippocampus because this is the most epileptogenic zone in patients and animal models (Buckmaster et al., 2022; Wyeth et al., 2020). Zbtb14 immunostaining was observed in the cytoplasm and cell nuclei. Moreover, the Zbtb14 protein-expressing cells were also NeuN-positive, indicating neuronal localization of the Zbtb14 expression (Figure 1).

**Figure 1.**
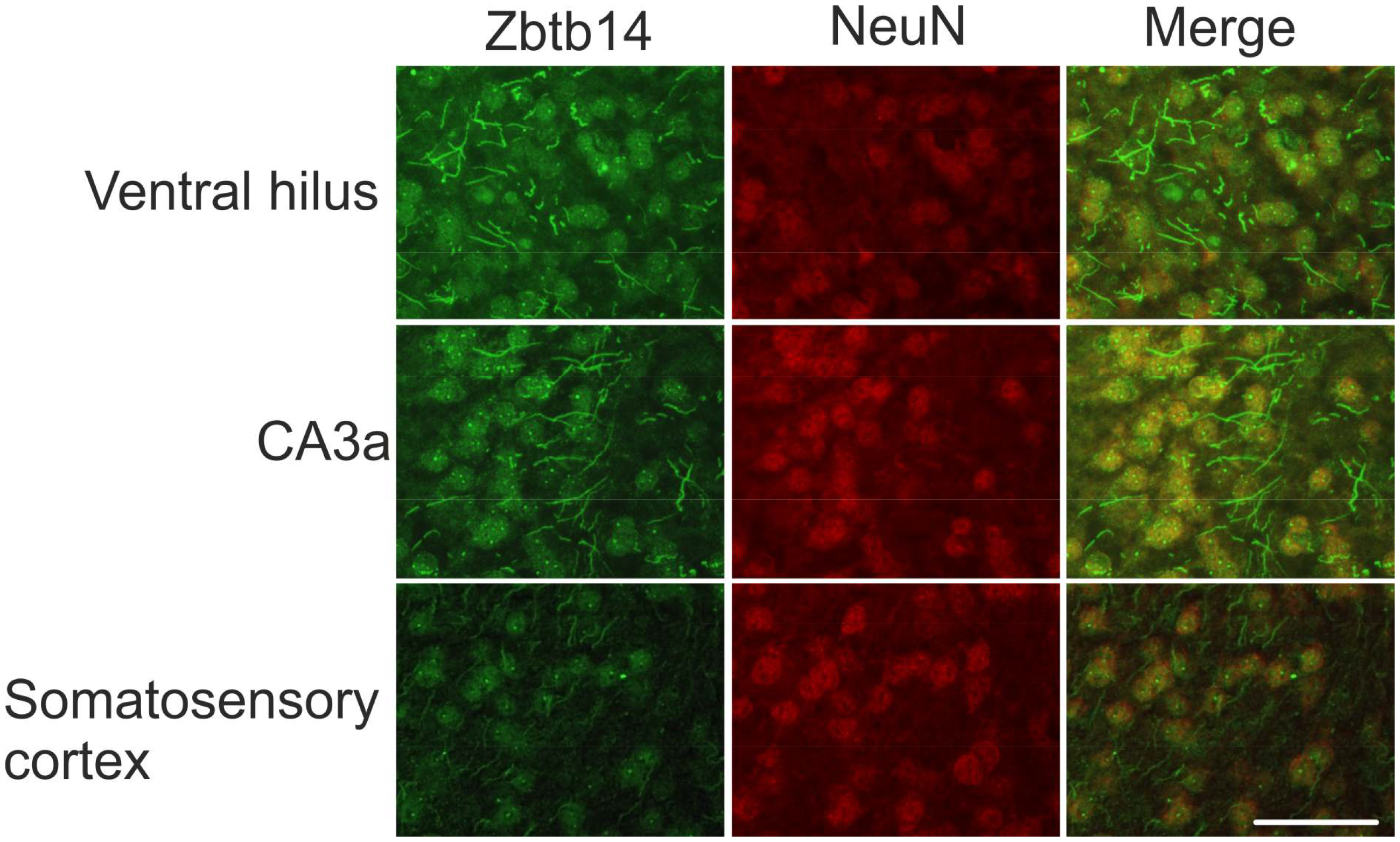
Cellular localization of Zbtb14 and NeuN at 3 AM in the hippocampus and the somatosensory cortex. Representative images from the ventral hilus, CA3a of the hippocampus, and layer VI of the somatosensory cortex of naive mice. Scale bar: 50 μm.

### The circadian dynamics of Zbtb14 protein levels in the cytoplasmic and nuclear compartments

Zbtb14 protein levels through the circadian cycle were evaluated separately in the cytoplasmic and nuclear extracts. Data are presented as multiples of the first time point after the light was turned on, i.e., 11 AM (Figure 2). In the cytoplasm, the Zbtb14 protein level increased by 1.45±0.05 at 3 PM compared to 11 AM (p<0.05) and then gradually decreased to 1.13±0.11 at 7 PM, 0.84±0.08 at 11 PM (p<0.01 compared to 3 PM), 0.73±0.11 at 3 AM (p<0.001 compared to 3 PM), and 0.54±0.16 at 7 AM (p<0.05 compared to 11 AM; p<0.001 compared to 3 PM, and p<0.01 compared to 7 PM). No significant differences were observed in the Zbtb14 protein level in the cytoplasm during the light-off phase.

**Figure 2.**
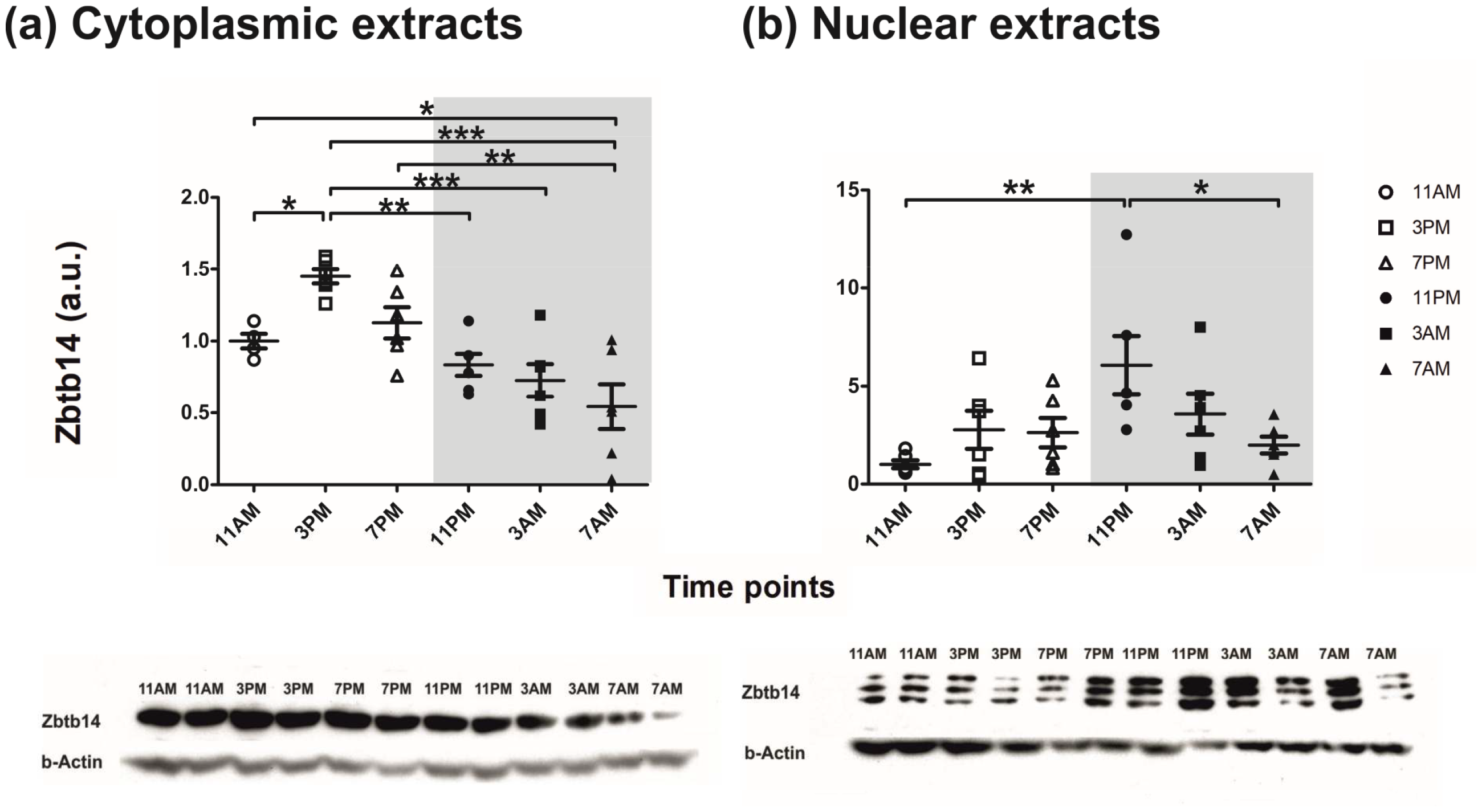
Relative levels of Zbtb14 protein in the cytoplasm and nuclei in the hippocampus throughout the circadian cycle. The data are expressed as multiples of the relative ratios at 11 AM in the cytoplasmic **(a)** or nuclear **(b)** extracts, respectively. The relative ratios of Zbtb14 to β-actin are represented as the mean ± SEM (n=6, *p<0.05, **p<0.01, ***p<0.001, one-way ANOVA with Tukey’s multiple comparison post hoc test).

Different dynamics were observed in Zbtb14 protein levels in the nuclear extracts. During the light-on phase, the level of Zbtb14 was not different from the level at 11 AM and was 2.76±0.96 at 3 PM, and 2.61±0.75 fold greater at 7 PM. At 11 PM, the level of Zbtb14 was 6.07±1.48 fold greater than at 11 AM (p<0.01). Then, the levels gradually decreased to 3.57±1.05 at 3 AM and then to 1.98±0.43 at 7 AM, significantly lower than at 11 PM (p<0.05).

### The circadian dynamics of the density of Zbtb14-expressing cells

We then assessed the temporal pattern of the density of Zbtb14-expressing cells in the dorsal and ventral hilus, dorsal and ventral CA3a of the hippocampus, and individual layers of the somatosensory cortex using immunostaining (Figure 3). Representative images of the Zbtb14 staining pattern at 3 PM, 11 PM, and 3 AM are presented in Figure 4.

**Figure 3.**
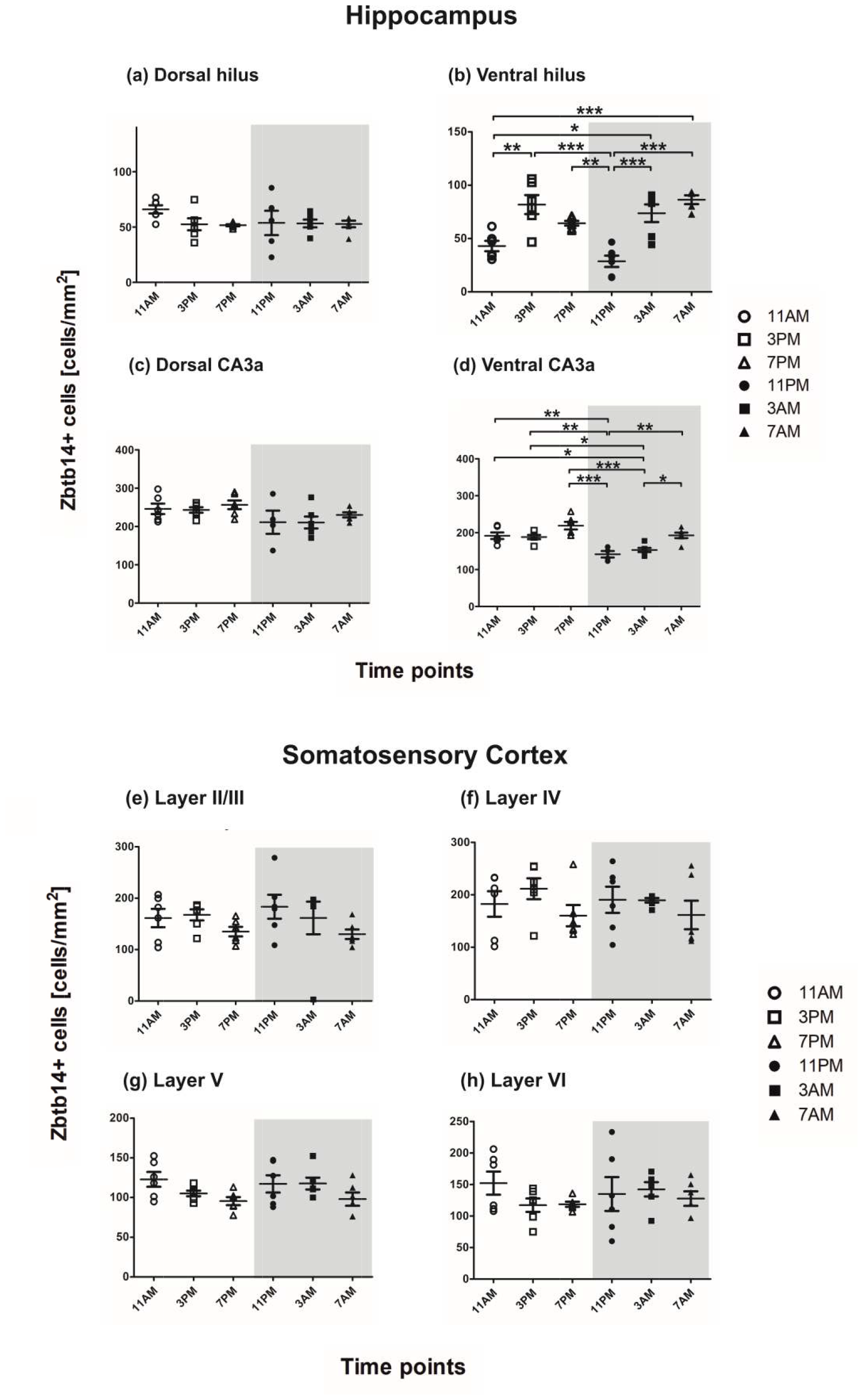
Zbtb14 protein-positive cell densities in the dorsal and ventral hilus and CA3a of the hippocampus and the somatosensory cortex. The density of Zbtb14-positive cells was counted in **(a)** dorsal hilus (n=5 for the time-point 11 PM, and n=6 for the other time points), **(b)** ventral hilus (n=5 for the time-point 7 AM, and n=6 for the other time points), **(c)** dorsal CA3a (n=4 for the time-point 11 PM, and n=6 for the other time points), **(d)** ventral CA3a (n=4 for the time-point 11 PM, and n=6 for the other time points), **(e)** layer II/ III (n=6 per time-point), **(f)** layer IV (n=6 per time-point), **(g)** layer V (n=6 per time-point), and **(h)** layer VI (n=6 per time-point). Values are represented as the mean number of cells/mm^2^ ± SEM. Data are analyzed by one-way ANOVA with Tukey’s multiple comparison post hoc test (*p<0.05, **p<0.01, ***p<0.001).

**Figure 4.**
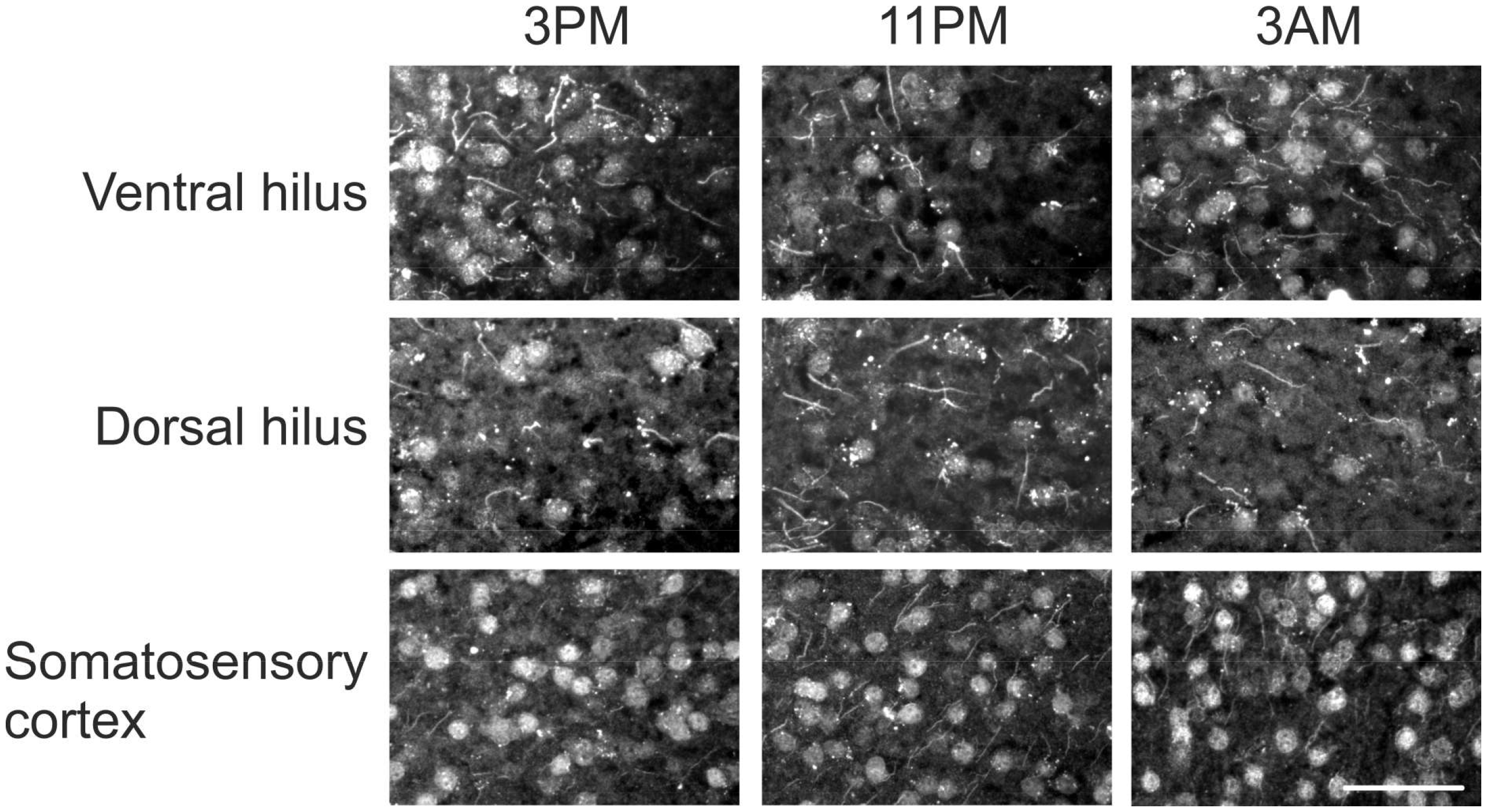
Zbtb14 immunostaining at 3 PM, 11 PM, and 3 AM in the dorsal and ventral hilus and layer VI of the somatosensory cortex. Representative images from 3 PM, 11 PM, and 3 AM time points in the ventral and dorsal hippocampal hilus and the somatosensory cortex. Note the lower expression of Zbtb14 in the ventral hilus at 11 PM—scale bar: 50 μm.

The density of Zbtb14-positive cells showed no significant change at the observed time points in the dorsal hilus, dorsal CA3a, and the somatosensory cortex (Figure 3 a, c, e-h). On the contrary, the density of Zbtb14-positive cells revealed significant differences throughout the circadian cycle in the ventral hilus and CA3a areas (Figure 3 b and d). The mean density of Zbtb14-positive cells in the ventral hilus at 11 AM, the first time point after the light was on, was 43.01±4.88 cells/mm^2^. It increased significantly at 3 PM to 81.85±8.88 cells/mm^2^ (p<0.01 compared to 11 AM). Then, it decreased steadily, reaching 64.35±2.22 cells/mm^2^ at 7 PM and 28.68±5.31 cells/mm^2^ at 11 PM (p<0.001 compared to 3 PM and p<0.01 compared to 7 PM). Next, the density of Zbtb14-positive cells increased again and was 73.74±8.25 cells/mm^2^ at 3 AM (p<0.001 compared to 11 PM; p<0.05 compared to 11 AM, and 86.40±4.14 at 7 AM (p<0.001 compared to 11 PM; p<0.001 higher compared to 11 AM).

The mean density of Zbtb14-positive cells in the ventral CA3a at 11 AM was 191.6±8.91 cells/mm^2^ and remained unchanged at 3 PM, 188.1±5.73 cells/mm^2^, and at 7 PM, 218.9 ±10.26 cells/mm^2^. During the light-off phase, Zbtb14-positive cell density first dropped down to 141.4±8.83 cells/mm^2^ at 11 PM (p<0.001 compared to 7 PM, p<0.01 compared to 11 AM and 3 PM) and then started to increase to 153.0± 5.65 cells/mm^2^ at 3 AM (p<0.001 compared to 7 PM, p<0.05 compared to 11 AM and 3 PM), and to 192.7±7.47 cells/mm^2^ at 7 AM (p<0.01 compared to 11 PM, p<0.05 compared to 3 AM), which was not different from the density at 11 AM.

The overall data suggest that circadian rhythm modulates Zbtb14 protein expression in the ventral hippocampus but not in the dorsal and somatosensory cortex.

### Overrepresentation of ZF5 motif in promoters of genes downregulated in the in vitro model of epileptiform discharges

A transcriptomics data set containing gene expression profiles 24 hours after induction of epileptiform discharges in neurons *in vitro* was used to evaluate the frequency of the ZF5 motif in genes whose expression was altered. *In silico* analysis revealed the overrepresentation of the ZF5 motif in the promoters of genes affected. The ZF5 motif was present only in promoters of down-regulated genes and was detected in promoters of 21 out of 24 down-regulated genes, which is significantly different (p_adj_ = 7.233x 10^−4^) than expected (Table 1).

**Table 1.**
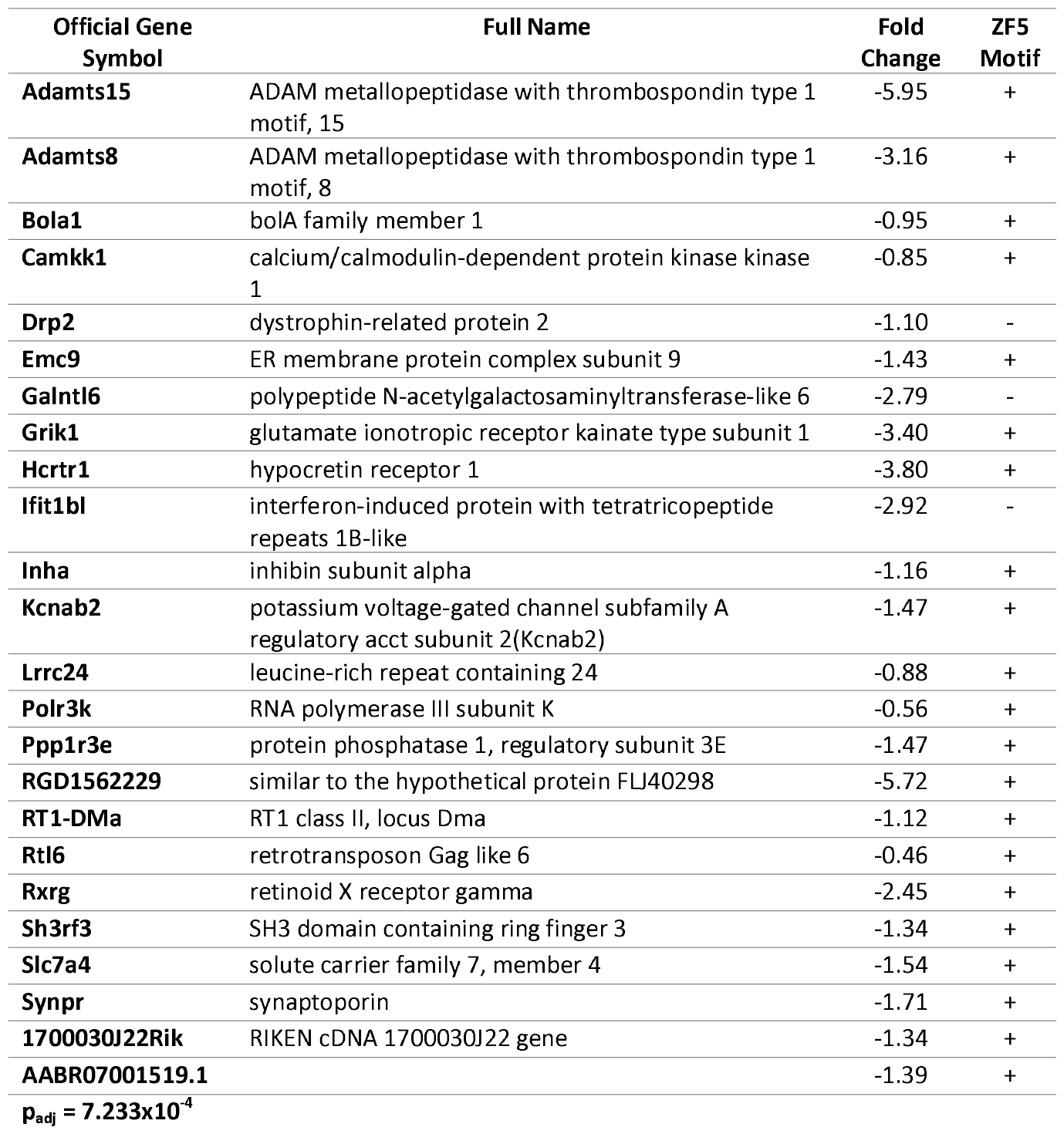
The presence of the ZF5 motif (GSGCGCGR; TF: M00716_1) in the promoters of genes downregulated in an *in vitro* model of epileptiform discharges.

### Zbtb14 protein expression in an in vivo model of a temporal lobe epilepsy

The expression of Zbtb14 in the hippocampus was evaluated in the pilocarpine model of epilepsy in mice. Tissues were collected at two selected time points: 3 PM and 11 PM (Figure 5). No difference was observed in protein levels between control and epileptic animals at 3 PM (1.64±0.62 vs. 2.84±0.61 a.u., respectively). However, at 11 PM, the expression of Zbtb14 was significantly higher in epileptic animals than in controls (19.95±2.24 vs. 3.73 ±1.16 a.u. respectively, p<0.001). Expression of Zbtb14 in epileptic animals was significantly higher at 11 PM as compared to 3 PM (p<0.001).

**Figure 5.**
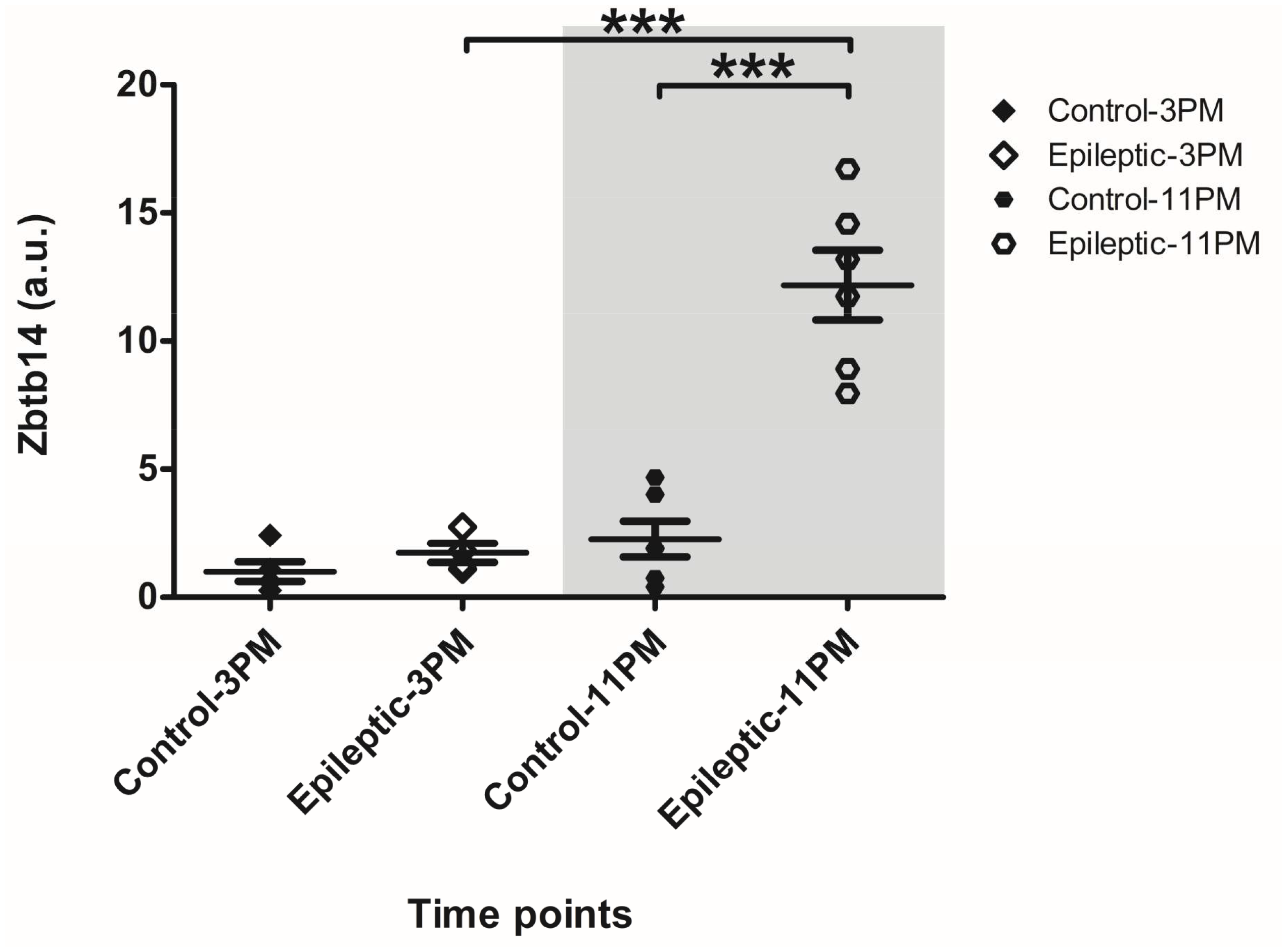
Zbtb14 protein expression in the hippocampus in pilocarpine-induced epilepsy. The relative ratios of Zbtb14 to β-actin are represented as the mean ± SEM (n=5 for Control-3 PM, n=4 for Epileptic-3 PM, and n=6 for Control- and Epileptic-11 PM; *p<0.05, **p<0.01, ***p<0.001, one-way ANOVA with Bonferroni’s multiple comparison post hoc test).

## Discussion

Our work revealed that: (i) the Zbtb14 protein is expressed in neurons in the mouse brain; (ii) Zbtb14 protein levels oscillate through the circadian cycle in the ventral hippocampus but not in the dorsal hippocampus; (iii) the oscillation of the Zbtb14 protein occurs in both the cytoplasm and nucleus but in a different temporal pattern; (iv) the circadian dynamics of the Zbtb14 protein are perturbed in epilepsy in an *in vivo* model of epilepsy; (v) numerous genes that are downregulated in the *in vitro* model of epileptiform discharges have a ZF5 motif in their promoters.

Overrepresentation of the ZF5 motif in the promoters of clock-controlled genes in the heart, liver, muscles, suprachiasmatic nucleus, and the ventral hippocampus has already been suggested based on *in silico* analysis (Bozek et al., 2009; Debski et al., 2020). The notion that densities of Zbtb14-positive cells oscillate during the circadian cycle only in the ventral and not in the dorsal hippocampus is attractive regarding the hippocampal heterogeneity over the longitudinal axis. The ventral hippocampus differs in connectivity, gene expression pattern, neurochemical pattern, and function from the dorsal hippocampus in rodents (Bienkowski et al., 2018; Brancati et al., 2021; Lothmann et al., 2021; O’Leary and Cryan, 2014). The oscillation of Zbtb14, and potentially its targets, in the ventral hippocampus may play a role in the circadian regulation of the functions, which are specific to the ventral hippocampus, a hypothesis that remains to be tested.

The ventral hippocampus is a primary epileptogenic zone in pilocarpine-induced epilepsy in rodents and also in the anterior hippocampus in humans (Buckmaster et al., 2022; Wyeth et al., 2020). Interestingly, we have found that the circadian dynamics of Zbtb14 are altered in epileptic animals. At 11 pm, epileptic animals had almost six times higher levels of Zbtb14 protein in whole tissue extracts than control animals. The 11 pm time-point is the peak expression time-point of the Zbtb14 protein in nuclear extracts. The higher expression of the Zbtb14 protein may be related to the higher accessibility of Zbtb14 in the nucleus for transcriptional regulation. This would presumably result in the altered expression of the Zbtb14 target genes.

The analysis of the effect of the epileptiform discharges *in vitro* revealed that 21 out of 24 downregulated genes contain the ZF5 motif in their promoters. Interestingly, 11 of those are expressed in the mouse brain: Adamts15, Bola1, Camkk1, Grik1, Kcnab2, Lrrc24, Ppp1r3e, Sh3rf3, Slc7a4, Synpr, and 1700030J22Rik according to the Allen Brain Atlas. An additional six genes were found to be detected in the mammalian brain according to a PubMed search: Adamts8 (Dunn et al., 2006; Rossier et al., 2015) Hcrtr1 (Li et al., 2018; Scott et al., 2011), Inha (Fujimura et al., 1999), Polr3k (Lata et al., 2021), Rtl6 (Irie et al., 2022), and Rxrg (McCullough et al., 2018). Many of the above genes have essential roles in the brain function. For example, Adamts15 and Adamts8 are metalloproteases that reshape perineuronal nets (Rossier et al., 2015); Kcnab2 is a subunit of voltage-gated potassium channel complexes that play an essential part in neuronal excitability and is a risk factor for epilepsy (Kurosawa et al., 2005; Yee et al., 2022); Synpr is a component of synaptic vesicle membranes and the marker of sprouting of the mossy fiber in the hippocampus (Cabrera et al., 2022; Knaus et al., 1990); Grik1 encodes an ionotropic glutamate receptor subunit known as GluK1 and is involved in synaptic transmission and plasticity (Valbuena et al., 2019); Hcrtr1, hypocretin/orexin receptor 1 can induce neurotransmitter release both post- and pre-synaptically, is known to play a role in the sleep-wake cycle, seizures and anxiety (Kordi Jaz et al., 2017; Li et al., 2018; Scott et al., 2011); Retinoid X receptor gamma (Rxrg) is part of a nuclear receptor family and is activated by retinoic acid supports the survival of dopaminergic neurons (Friling et al., 2009); Polr3k is one of the subunits RNA polymerase III which decreased activity perturbs protein and neurotransmitter shuttling via the ER and affects synaptic plasticity in both axons and dendrites (Lata et al., 2021). The Zbtb14 protein seemingly regulates the genes associated with synaptic activity, transmission, and plasticity.

Our previous transcriptomics data (Debski et al., 2020) from the ventral hippocampus of control and epileptic animals share three differentially expressed genes with the *in vitro* data presented in this paper: Adamts15, 1700030J22Rik, and Inha. Debski et al. reported that *in vivo* Adamts15 and 1700030J22Rik did not have circadian oscillations in control animals but gained oscillatory patterns in pilocarpine-treated epileptic animals (Debski et al., 2020). This circadian dynamics change in Adamts15 might result in altered regulatory function on the extracellular matrix of Adamts15 that is triggered by epilepsy. Conversely, Inha oscillates throughout the circadian cycle in control animals, but this rhythm is abolished in epileptic animals. The overlap of three genes, Adamts15, 1700030J22Rik, and Inha, from two different experimental datasets, strengthens the possibility of the involvement of the Zbtb14 transcription factor in their regulation.

Although epileptic seizures appear as random events, clinical data, patient journals, and basic research have shown that seizures occur more often at certain times over the circadian cycle (Hofstra et al., 2009; Karoly et al., 2021; Sun and Wang, 2023; Zhang et al., 2021). The investigation of cycles from circadian to circannual is becoming an important research topic due to the prospect of chronotherapy – the administration of drugs when they are most effective. Whether the temporal pattern of seizure rhythmicity stems, at least in part, of proteins like Zbtb14 is an interesting hypothesis to test, as it may provide an interesting novel therapeutic target.

### Limitations of the study

In this study, we only used male animals to reduce the complexity inherent to accounting for the female estrous cycle. However, we recognize the need to include female animals in future studies as several epidemiological studies are emphasizing the sexual dimorphism in epilepsy, including the differences in the types of epilepsy females tend to have, seizure threshold, and epileptogenesis (Hophing et al., 2022; Reddy et al., 2021).

## Author Contributions

İlke Güntan – investigation, data analysis, writing; Antoine Ghestem – investigation; Kinga Nazaruk – investigation, data analysis; Karolina Nizińska – investigation; data analysis; Maciej Olszewski – investigation; Dorota Nowicka – investigation, supervision; Christophe Bernard – conceptualization, supervision, writing; Katarzyna Łukasiuk – conceptualization, supervision, funding acquisition, writing

## Data Availability

RNA seq data are deposited at the NCBI GEO repository with the submission number: GSE227084

